# A conserved role of Hippo signaling in initiation of the first lineage specification event across mammals

**DOI:** 10.1101/2022.07.01.498418

**Authors:** Claudia Gerri, Afshan McCarthy, Gwen Mei Scott, Marius Regin, Sophie Brumm, Claire S. Simon, Janet Lee, Cristina Montesinos, Caroline Hassitt, Sarah Hockenhull, Daniel Hampshire, Kay Elder, Phil Snell, Leila Christie, Ali A. Fouladi-Nashta, Hilde Van de Velde, Kathy K. Niakan

## Abstract

Our understanding of the molecular events driving cell specification in early mammalian development relies mainly on mouse studies, and it remains unclear whether these mechanisms are conserved across mammals, including humans. We have recently shown that the establishment of cell polarity via aPKC is a conserved event in the initiation of the trophectoderm (TE) placental program in mouse, cow, and human embryos. However, the molecular mechanisms transducing cell polarity into cell fate in cow and human embryos is unknown. Here, we have examined the evolutionary conservation of the molecular cascade downstream of aPKC in four different mammalian species: mouse, rat, cow, and human. Surprisingly, by morphokinetic and immunofluorescence analyses, we observe that rat embryos more closely recapitulate human and cow developmental dynamics, in comparison to the mouse. Nevertheless, in all four species, inhibition of the Hippo pathway by targeting LATS kinases is sufficient to drive ectopic TE initiation and downregulation of SOX2, a marker of the inner cell mass. Our comparative embryology approach uncovered intriguing differences as well as similarities in a fundamental developmental process among mammals, reinforcing the importance of cross-species investigations.

## Introduction

Our current understanding of the molecular mechanisms regulating cell specification during preimplantation mammalian development relies mainly on the mouse, which is a tractable model for molecular genetic analysis. Prior to implantation, at the 8-cell stage, the mouse embryo undergoes a drastic morphological change, whereby blastomeres flatten and adhere to each other in a process known as compaction (Ducibella and Anderson, 1975). After several rounds of cell division, two distinct cell populations are discernible at the morula stage: inner and outer cells. Upon reaching the blastocyst stage, inner cells give rise to the inner cell mass (ICM), and the outer cells become the trophectoderm (TE), a polarized epithelium that will form fetal components of the placenta. Subsequently, the ICM further segregates into the epiblast (EPI), which gives rise to the fetus, and the primitive endoderm (PrE), which primarily contributes to the yolk sac (Cockburn and Rossant, 2010).

Concomitant with compaction, cell polarity is established in the 8-cell mouse embryo. Inner and outer cells display different polarization states, which influence their cell fate acquisition. The contact-free surface of the outer cells acquires an apical domain, enriched with the atypical protein kinase C (aPKC) that together with the proteins partitioning defective homolog 6B (PARD6B) and homolog 3 (PARD3), forms the PAR complex which specifies the apical domain, while PAR1 (EMK1 or MARK2), E-CADHERIN and other cell adhesion molecules localize to the basolateral domain (Plusa et al., 2005, Vinot et al., 2005, Ohsugi et al., 1996, Alarcon, 2010). In the polar outer cells, the apical PAR proteins sequester Angiomotin (AMOT), a modulator of the Hippo pathway, from the junctional complexes. This interaction prevents activation of downstream Hippo pathway kinases, Large tumor suppressor kinases 1/2 (LATS1/2) (Hirate et al., 2013). Consequently, in outer cells, Yes-associated protein 1 (YAP1) and WW domain-containing transcription regulator protein 1 (WWTR1, also known as TAZ) accumulate in the nucleus where they bind the TEA-domain family member 4 (TEAD4), to promote the expression of TE lineage-associated factors, such as *Caudal type homeobox 2 (Cdx2)* and *Gata binding protein 3 (Gata3)* (Strumpf et al., 2005, Ralston et al., 2010, Nishioka et al., 2009). By contrast, in the apolar inner cells, AMOT is free to interact and is activated through phosphorylation by LATS1/2. In these cells, activation of the Hippo pathway results in YAP1 and WWTR1 phosphorylation and cytoplasmic retention, thus maintaining the inner cells in an unspecified state (Hirate et al., 2013, Cockburn et al., 2013, Frum et al., 2018, Leung and Zernicka-Goetz, 2013). SOX2 is a molecular marker of the epiblast and considered to be the earliest in its restriction to inner cells at the morula stage in mice (Wicklow et al., 2014). SOX2 expression has been shown to be restricted to inner cells by repression of YAP1/WWTR1/TEAD4 in the outer cells (Frum et al., 2018, Frum et al., 2019). These mechanistic insights derive from functional studies performed in the mouse embryo, and it is unclear to what extent these phenomena are conserved in other mammals.

In the mouse, CDX2 is more strongly expressed in outer cells from the morula stage (Dietrich and Hiiragi, 2007), and *Cdx2* mutant embryos exhibit loss of epithelial integrity at the blastocyst stage, thus failing to maintain the blastocoel cavity or to implant (Strumpf et al., 2005). Notably, in human and cow embryos, CDX2 is detectable only at a later stage, in cavitating blastocysts (Niakan and Eggan, 2013, Goissis and Cibelli, 2014a, Berg et al., 2011). Moreover, *Cdx2* knockdown experiments in cow embryos demonstrated that CDX2 is required only later in TE maintenance and does not repress *Oct4* expression (Berg et al., 2011), as it has been suggested in mouse (Niwa et al., 2005). Cow embryos with impaired CDX2 activity appear phenotyipicallly different to mouse embryos and it has been proposed that the regulatory circuitry involved in TE/ICM specification is fundamentally different in these two species (Berg et al., 2011). Altogether, these findings hint at interesting differences in the molecular cascade controlling cell specification. We have previously shown that morula stage mouse, cow, and human embryos, acquire an apical–basal cell polarity in outer cells, with expression of aPKC localized to the contact-free domain, nuclear expression of the Hippo pathway downstream effectors YAP1 and WWTR1, and restricted expression of TE-associated factors such as GATA3, suggesting conservation of the initiation of a TE program across these species (Gerri et al., 2020). Furthermore, inhibition of aPKC activity impairs TE initiation at the morula stage in cow and human embryos (Gerri et al., 2020), and downregulation of the polarity regulators Phospholipase CB1 (PLC) and PLCE1 leads to decreased GATA3 expression in human embryos (Zhu et al., 2021).

In the present study, we expanded our morphokinetic and molecular analysis to another mammalian species, the rat. To our surprise, we found that the expression dynamics of molecular markers of inner and outer cells in the rat morula are more similar to cow and human than its closely related rodent species, the mouse. As a result of this, we set out to understand the role and evolutionary conservation of the molecular cascade driving TE fate downstream of aPKC, focusing on Hippo signaling pathway kinases. We found that in all the four species, inhibition of the LATS kinases promotes TE initiation in all cells, leading to ectopic YAP1 and GATA3 expression in the inner cells at morula stage and in the ICM of blastocysts, accompanied by a reduction of SOX2 expression. Interestingly, LATS inhibition completely depleted SOX2 expression in mouse embryos; however, in the other species, SOX2 expression was downregulated, but not completely abrogated. We observed intriguing differences in the expression pattern of SOX2 when comparing the mouse to the other mammalian species in physiological conditions, whereby SOX2 expression is restricted to inner cells in the mouse morula, but SOX2 is broadly expressed in all cell types in the other species. Altogether, these data indicate that while there is a conserved role for LATS kinases in preventing ectopic induction of TE in inner cells that go on to form the ICM, there are interesting species-specific differences in the regulation of the pluripotency factor SOX2 that warrants further investigation.

## Results and discussion

We initially performed morphokinetic analysis of rat embryos using similar criteria to our previous study (Gerri et al., 2020) (**Videos 1-4**). We calculated the duration of each morphological milestone as a percentage of time from the 8-cell stage to the end of cavitation. Mouse embryos remain at the 8-cell stage for a comparatively short period of time and rapidly undergo compaction (Gerri et al., 2020). By contrast, we observed an extended 8-cell-to-compaction transition with multiple cell divisions in rat, cow, human embryos (**Figure supplement 1**) (Gerri et al., 2020). We termed this stage as “pre-compaction”. In addition, mouse embryos exhibit a comparatively longer morula stage, while cow, rat, and human embryos show a rapid transition from pre-compaction to cavitation (**Figure supplement 1**) (Gerri et al., 2020). These data provide a framework for comparative analysis of preimplantation development in different species, as well as revealing important morphokinetic differences between the two rodent species considered in this study.

These intriguing results prompted us to analyze the expression pattern of key differentiation and pluripotency factors in rat embryos. First, we looked at the expression of TE-associated factors, such as YAP1 and GATA3. We observed weak nuclear expression of YAP1 in all blastomeres at 8-cell stage rat embryos (**Figure 1a, b**), similar to mouse embryos (Nishioka et al., 2009). At the morula stage, YAP1 and GATA3 are expressed mainly in the nucleus of outer cells in rat embryos. The expression of these factors remains stronger and mostly restricted to TE cells in rat blastocyst stage embryos (**Figure 1a, b**). Overall, the expression dynamics of YAP1 and GATA3 in rat embryos is similar to what has been reported in mouse, cow, and human embryos (Gerri et al., 2020).

**Figure 1.**
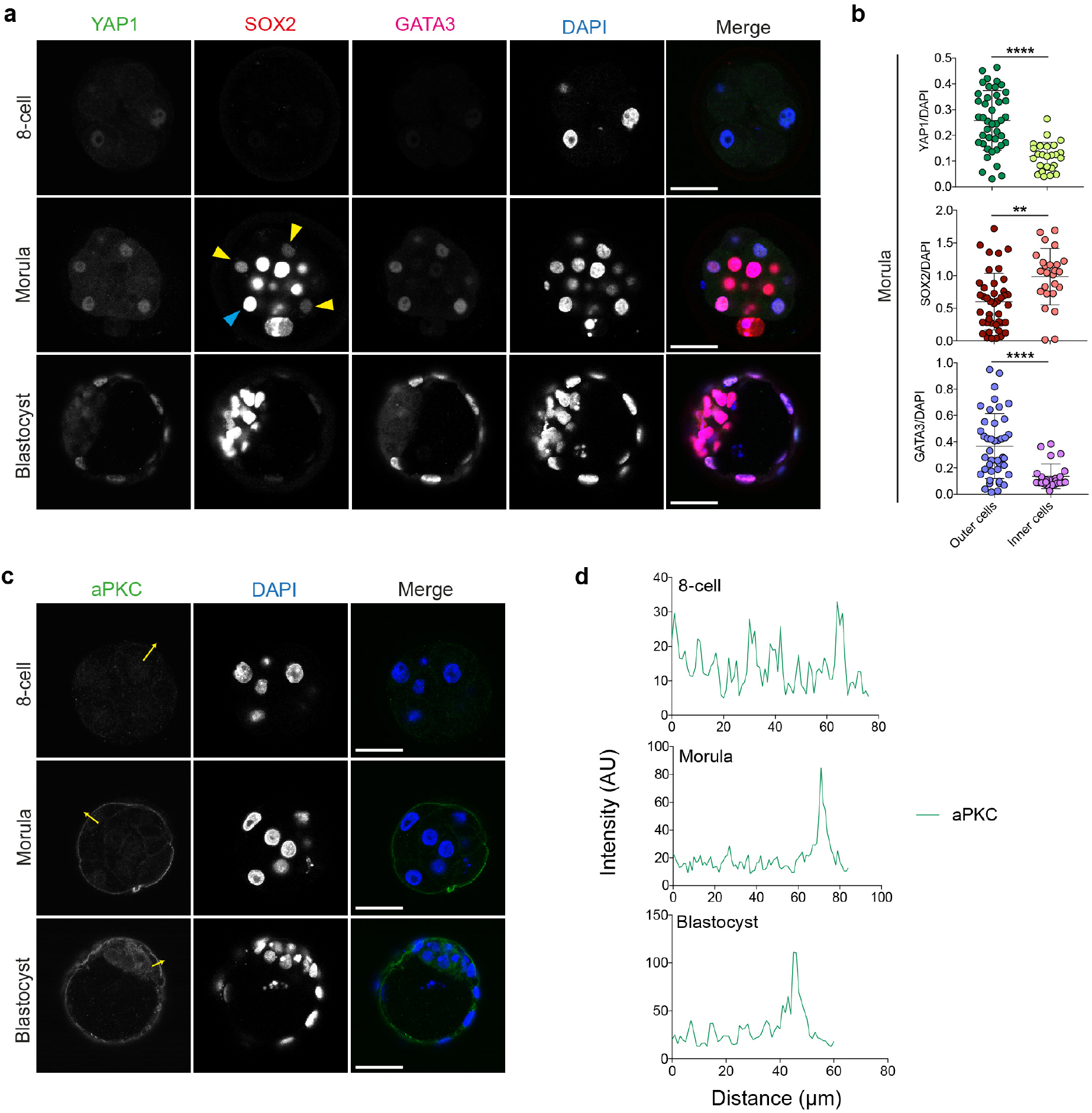
TE- and pluripotency-associated marker characterization in rat embryos. **a.** Time-course immunofluorescence analysis of YAP1, SOX2, GATA3 and DAPI nuclear staining in rat embryos at different developmental stage: 8-cell, morula and expanded blastocyst. Yellow arroheads point to outer cells with lower level of SOX2, while blue arrowhead shows an outer cell with high levels of SOX2. **b.** Quantification of YAP1, GATA3 and SOX2 normalized fluorescence intensity in rat morula stage embryos. (*n* = 67 cells from 5 embryos). Data are presented as mean ± s.d. Two-tailed Mann– Whitney *U* test, *\*\*\*P* < 0.001, \*\*\*\**P* < 0.0001. **c.** Time-course immunofluorescence analysis of aPKC and DAPI nuclear staining in rat embryos of different developmental stages: 8-cell, morula and expanded blastocyst. **d.** Fluorescence intensity profile of aPKC shown along the yellow arrows in the rat embryos shown in c. Scale bars, 30 um.

The transcription factor SOX2 is restricted in its expression to inner cells at the morula stage in mouse embryos and is considered the first marker of pluripotency in mouse inner cells (Wicklow et al., 2014). By contrast, SOX2 is expressed more broadly in both inner and outer cells in cow and human morula stage embryos, and is restricted to ICM cells only at later blastocyst stages (Gerri et al., 2020, Cauffman et al., 2009, Goissis and Cibelli, 2014b). Prompted by this striking difference between the mouse and other mammals, we assessed SOX2 expression dynamics in rat embryos. Surprisingly, SOX2 in rat morula stage embryos is detectable in both outer and inner cells and its expression is eventually restricted to the ICM cells only at blastocyst stage (**Figure 1a, b**), similar to human and cow embryos. Our previous work demonstrated regulation of YAP1 and GATA3 expression through the polarity complex in outer cells of morula stage embryos (Gerri et al., 2020). As rat embryos displayed a similar expression of these TE-associated transcription factors, we sought to investigate the expression of one of the upstream regulators of this pathway, aPKC, and observed apical localization in outer and TE cells at morula and blastocyst stage rat embryos (**Figure 1c, d**), similar to the other species (Gerri et al., 2020). Collectively, these data indicate conservation of aPKC protein localization and TE initiation at the morula stage in rat, cow, mouse and human embryos.

Downstream mechanism linking cell polarity and TE initiation in mammals other than the mouse remains unclear. To investigate molecular mechanisms downstream of aPKC activity we focused on the role of the Hippo pathway kinases LATS1/2. To functionally test the requirement of the LATS1/2 kinases in regulating early lineage initiation at the morula stage in rat, cow, and human embryos, we used a small molecule inhibitor known as TRULI, which has been characterized in mammalian tissues (Kastan et al., 2021). We first tested the inhibitor in mouse embryos, by treating embryos prior to compaction from 4-cell stage to the morula stage. We performed a dose-response test and found that the optimal concentration in mouse embryos, based on embryo viability and phenotypic assessment, was 5 μM (**Figure supplement 2 and Table supplement 1**). As expected from previous publications (Cockburn et al., 2013, Nishioka et al., 2009), we observed that LATS inhibition leads to significant ectopic expression of YAP1 and GATA3 in inner cells (**Figure 2a-c**). This is in contrast to DMSO volume-matched control embryos where YAP1 and GATA3 are exclusively localized to outer cells. Treated embryos also exhibited significant downregulation of SOX2 expression in the inner cells (**Figure 2a-c**), consistent with results following constitutive activation of YAP1 (Frum et al., 2018). In addition, by treating embryos from 4-cell stage to the blastocyst stage with the LATS inhibitor we observed nuclear expression of YAP1 and GATA3 in both TE and ICM cells, and a lack of SOX2 expression in ICM cells (**Figure 2a, d, e**). These results demonstrated that the LATS inhibitor phenocopies results from genetic modification of the Hippo signaling pathway in the mouse and therefore represents a useful method to investigate the role of the Hippo pathway in early mammalian embryos. Interestingly, we observed a significant increase in YAP1 and GATA3 expression in outer cells (**Figure 2c**), which may reflect residual LATS activity in these cells, which is blocked by LATS inhibitor treatment.

**Figure 2.**
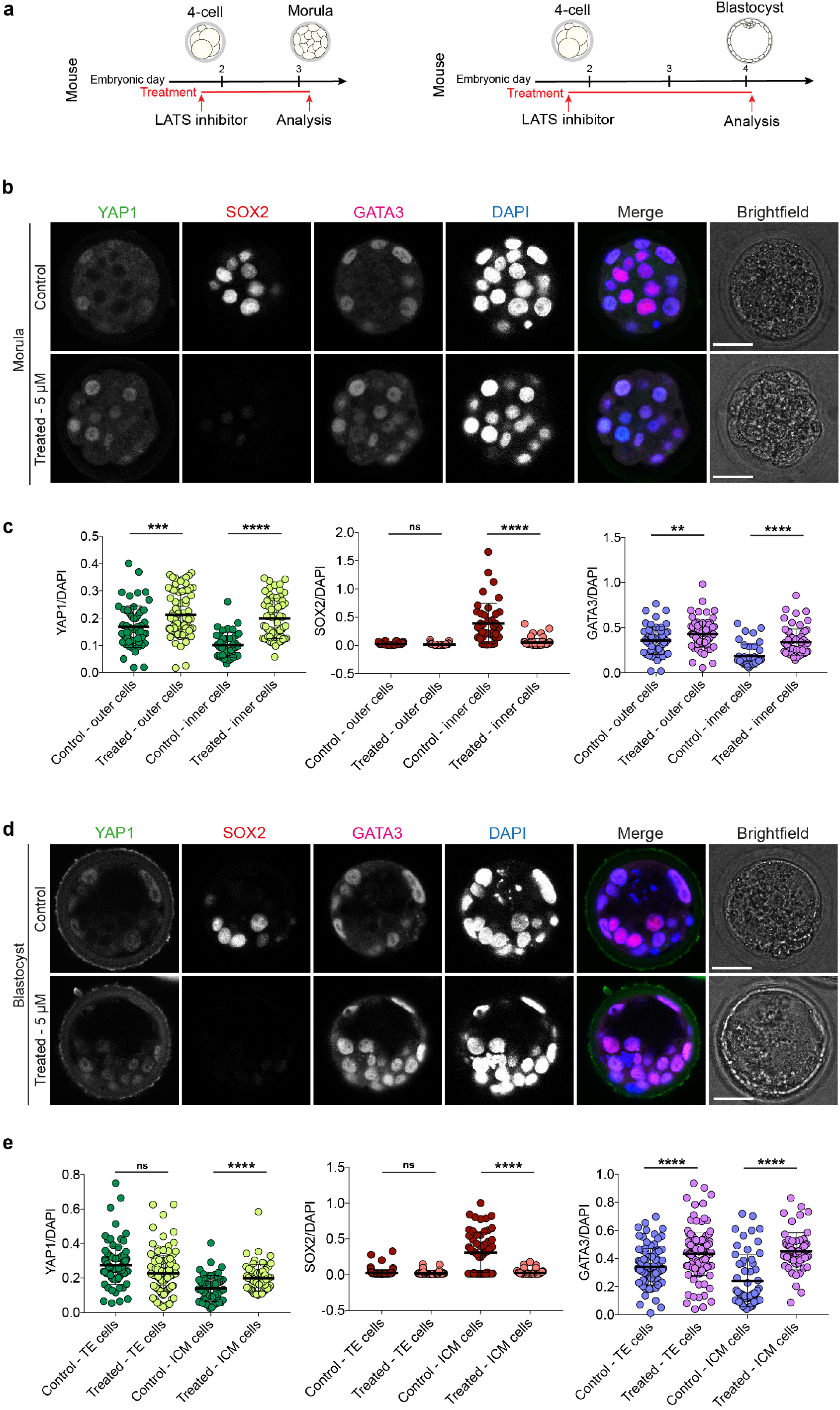
LATS inhibitor treatment in mouse embryos. **a.** Schematics of LATS inhibitor treatments in mouse embryos. **b.** Immunofluorescence analysis of YAP1, SOX2, GATA3 and DAPI nuclear staining in control and LATS-inhibitor-treated mouse morula stage embryos. **c.** Quantification of YAP1, SOX2, GATA3 normalized fluorescence intensity in outer and inner cells in control and LATS-inhibitor-treated mouse morula stage embryos. *(n* = 246 cells from 15 embryos). Data are presented as mean ± s.d. Two-tailed Mann–Whitney *U* test, ns = not significant, \*\**P* < 0.01, \*\*\**P* < 0.001, *\*\*\*\*P* < 0.0001. **d.** Immunofluorescence analysis of YAP1, SOX2, GATA3 and DAPI nuclear staining in control and LATS-inhibitor-treated mouse blastocyst stage embryos. **e.** Quantification of YAP1, SOX2, GATA3 normalized fluorescence intensity in TE and ICM cells in control and LATS-inhibitor-treated mouse blastocyst stage embryos. (*n* = 354 cells from 11 embryos). Data are presented as mean ± s.d. Two-tailed Mann–Whitney *U* test, ns = not significant, \*\*\*\**P* < 0.0001. Scale bars, 30 um.

We next extended our functional analysis to investigate the role of the LATS kinases in rat, cow, and human embryos. We initially performed dose response experiments in rat embryos and found that the optimal concentration of LATS inhibitor was 1 μM (**Figure supplement 3 and Table supplement 2**). Similarly to the mouse, LATS inhibitor-treated morula stage embryos exhibited significant ectopic YAP1 and GATA3 expression in inner cells, compared to control embryos (**Figure 3a-c**). SOX2 expression was significantly reduced in both outer and inner cells (**Figure 3a-c**); however, its expression was not entirely depleted, in contrast to mouse morula stage embryos where expression is largely undetectable (**Figure 2**). The expression of SOX2 in rat embryos treated with LATS inhibitor was abrogated only at blastocyst stage, with concomitant strong nuclear YAP1 and GATA3 in both TE and presumptive ICM cells (**Figure 3a, d, e**). These data suggest that, similar to the mouse, the Hippo pathway in rat embryos has a functional role in regulating YAP1 and GATA3 expression. Perdurance of SOX2 expression suggests that it might be indirectly regulated in rat embryos by mechanisms downstream of Hippo signaling kinases, in contrast to direct regulation of SOX2 by this pathway in the mouse (Frum et al., 2019), or that there may be differences in the timing of protein turnover. Moreover, while we observed a significant increase in YAP1 expression in outer cells, GATA3 expression was unaffected at the morula stage in rat embryos (**Figure 3c**), indicating that mouse embryos exhibit more rapid changes in the expression of these factors following inhibition of Hippo signaling pathway components.

**Figure 3.**
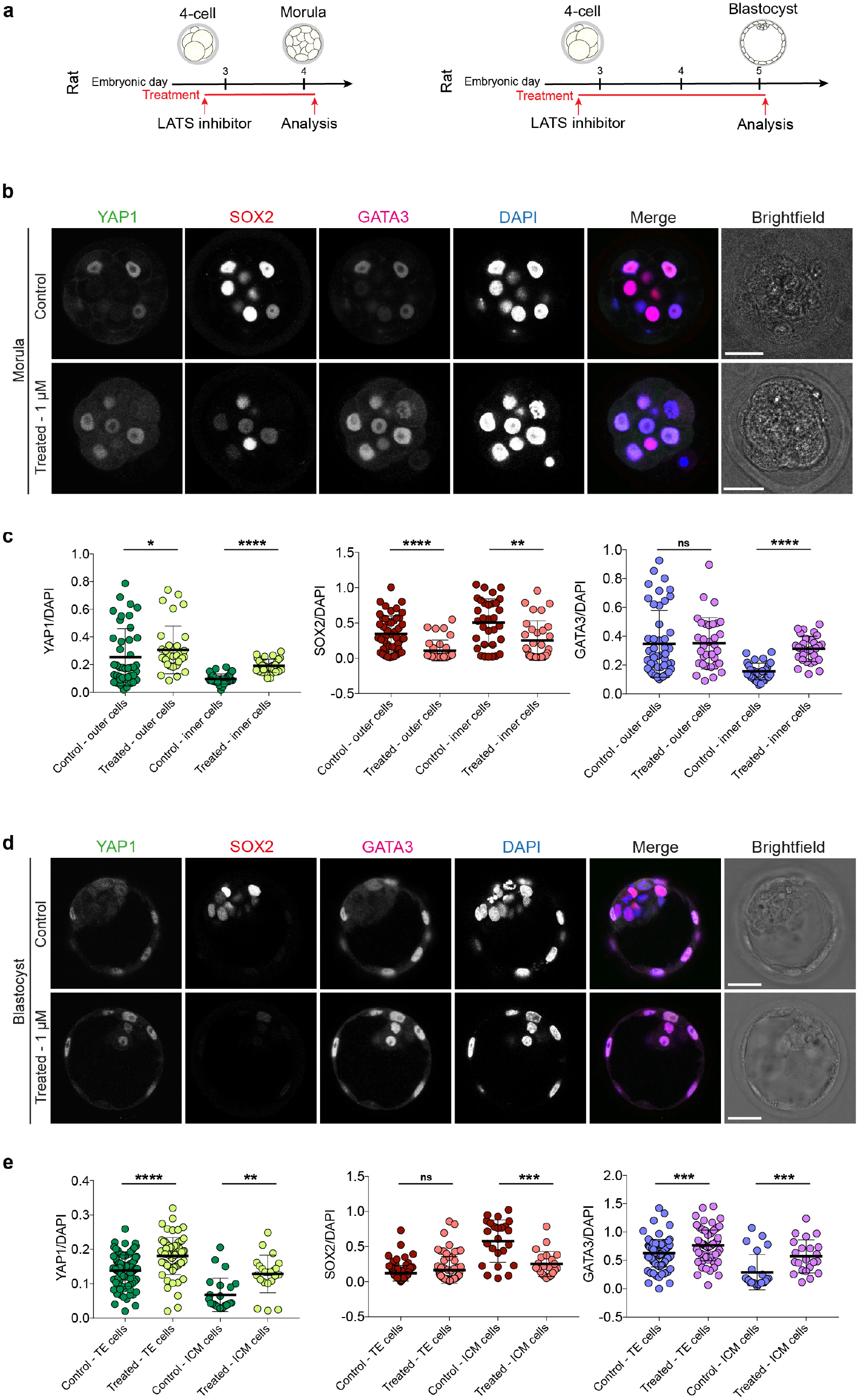
LATS inhibitor treatment in rat embryos. **a.** Schematics of LATS inhibitor treatments in rat embryos. **b.** Immunofluorescence analysis of YAP1, SOX2, GATA3 and DAPI nuclear staining in control and LATS-inhibitor-treated rat morula stage embryos. **c.** Quantification of YAP1, SOX2, GATA3 normalized fluorescence intensity in outer and inner cells in control and LATS-inhibitor-treated rat morula stage embryos. (*n* = 139 cells from 15 embryos). Data are presented as mean ± s.d. Two-tailed Mann–Whitney *U* test, ns = not significant, \**P* < 0.05, \*\**P* < 0.01, \*\*\*\**P* < 0.0001. **d.** Immunofluorescence analysis of YAP1, SOX2, GATA3 and DAPI nuclear staining in control and LATS-inhibitor-treated rat blastocyst stage embryos. **e.** Quantification of YAP1, SOX2, GATA3 normalized fluorescence intensity in TE and ICM cells in control and LATS-inhibitor-treated rat blastocyst stage embryos. (*n* = 207 cells from 12 embryos). Data are presented as mean ± s.d. Two-tailed Mann–Whitney *U* test, ns = not significant, \*\**P* < 0.01, \*\*\**P* < 0.001, \*\*\*\**P* < 0.0001. Scale bars, 30 um.

We next treated cow embryos with the LATS inhibitor. A dose response experiment showed that the optimal concentration of LATS inhibitor for cow embryos was 10 μM (**Figure supplement 4 and Table supplement 3**). Cow embryos treated from pre-to post-compaction with LATS inhibitor show ectopic expression of YAP1 and GATA3 in the nucleus of inner cells (**Figure 4a-c**). Upon LATS inhibition, SOX2 expression decreased but was not completely abrogated in both outer and inner cells at the morula stage (**Figure 4a-c**), an expression pattern similar to rat embryos (**Figure 3**). Longer treatment with the LATS inhibitor until the blastocyst stage resulted in sustained ectopic expression of YAP1 and GATA3 and a complete repression of SOX2 (**Figure 4a, d, e**).

**Figure 4.**
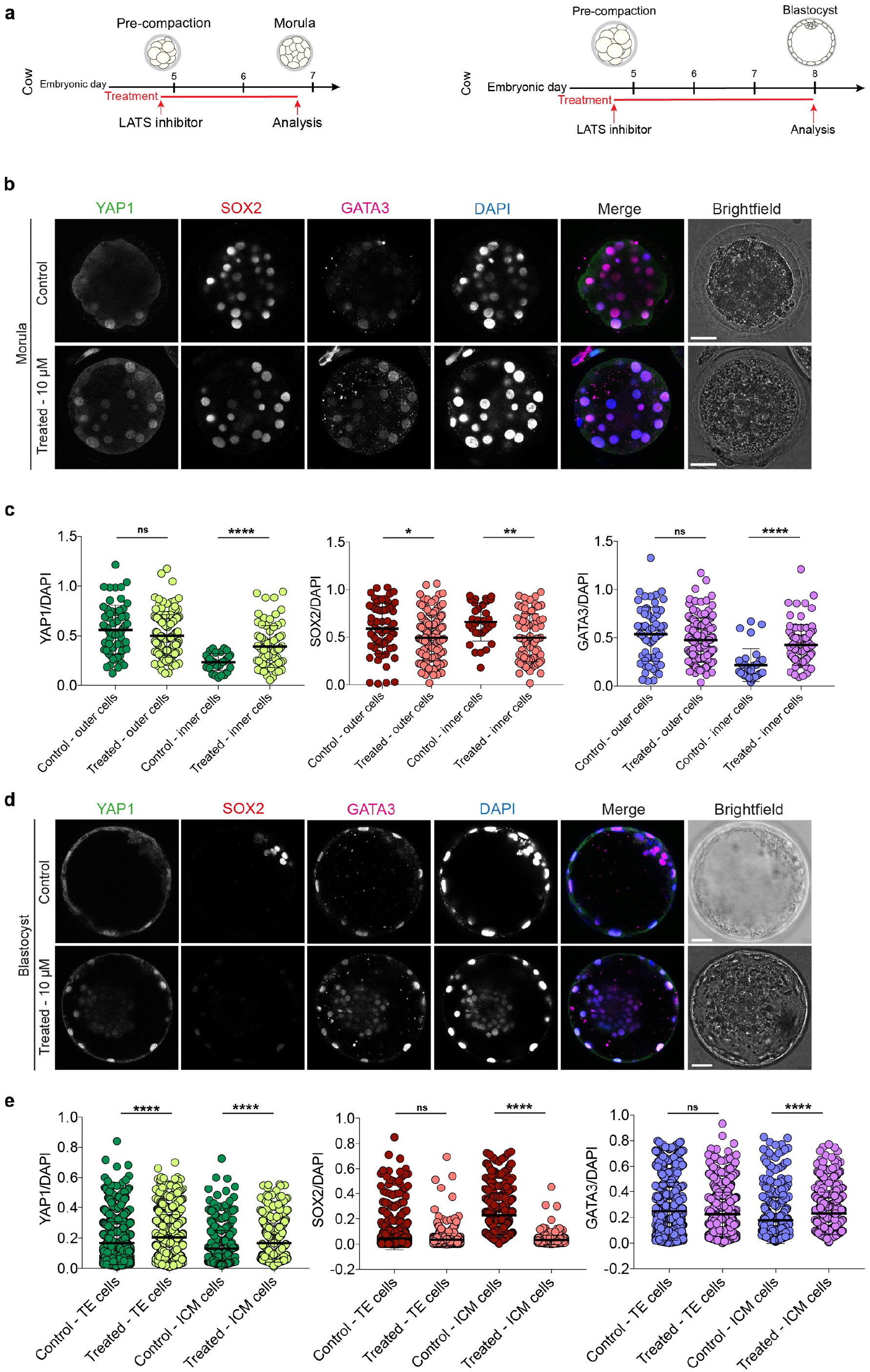
LATS inhibitor treatment in cow embryos. **a.** Schematics of LATS inhibitor treatments in cow embryos. **b.** Immunofluorescence analysis of YAP1, SOX2, GATA3 and DAPI nuclear staining in control and LATS-inhibitor-treated cow morula stage embryos. **c.** Quantification of YAP1, SOX2, GATA3 normalized fluorescence intensity in outer and inner cells in control and LATS-inhibitor-treated cow morula stage embryos. (*n* = 269 cells from 11 embryos). Data are presented as mean ± s.d. Two-tailed Mann–Whitney *U* test, ns = not significant, \*\**P* < 0.01, \*\*\*\**P* < 0.0001. **d.** Immunofluorescence analysis of YAP1, SOX2, GATA3 and DAPI nuclear staining in control and LATS-inhibitor-treated cow blastocyst stage embryos. **e.** Quantification of YAP1, SOX2, GATA3 normalized fluorescence intensity in TE and ICM cells in control and LATS-inhibitor-treated cow blastocyst stage embryos. (*n* = 2825 cells from 19 embryos). Data are presented as mean ± s.d. Two-tailed Mann–Whitney *U* test, ns = not significant, \*\**P* < 0.01, \*\*\**P* < 0.001, \*\*\*\**P* < 0.0001. Scale bars, 30 um.

To understand the functional role of the Hippo pathway in early human development, we treated human embryos with the LATS inhibitor from the pre-compaction until the post-compaction morula stage (**Figure 5a**). A dose response experiment showed that the optimal concentration of LATS inhibitor for human embryos was 5 μM (**Figure supplement 5 and Table supplement 4**). Upon LATS inhibition, the inner cells of human embryos exhibited ectopic YAP1 and GATA3 expression (**Figure 5b,c**). Similar to cow and rat embryos, SOX2 expression was reduced, but not completely abrogated, in inner cells at the morula stage following LATS inhibitor treatment compared to control embryos (**Figure 5b,c**). Moreover, in both cow and human embryos, YAP1 and GATA3 expression levels were unaffected in the outer cells at the morula stage (**Figures 4b,c and 5b,c**), unlike the mouse (**Figure 2**). Interestingly, in inner cells at the morula stage we observed that the absence of cytoplasmic retention of YAP1 is conserved in rat, cow, and human embryos (**Figures 3b, 4b, 5b**), while the mouse exhibits YAP1 cytoplasmic retention (**Figure 2b**). This suggests differences between species in the timing of expression or mode of action of effectors of the Hippo signalling pathway.

**Figure 5.**
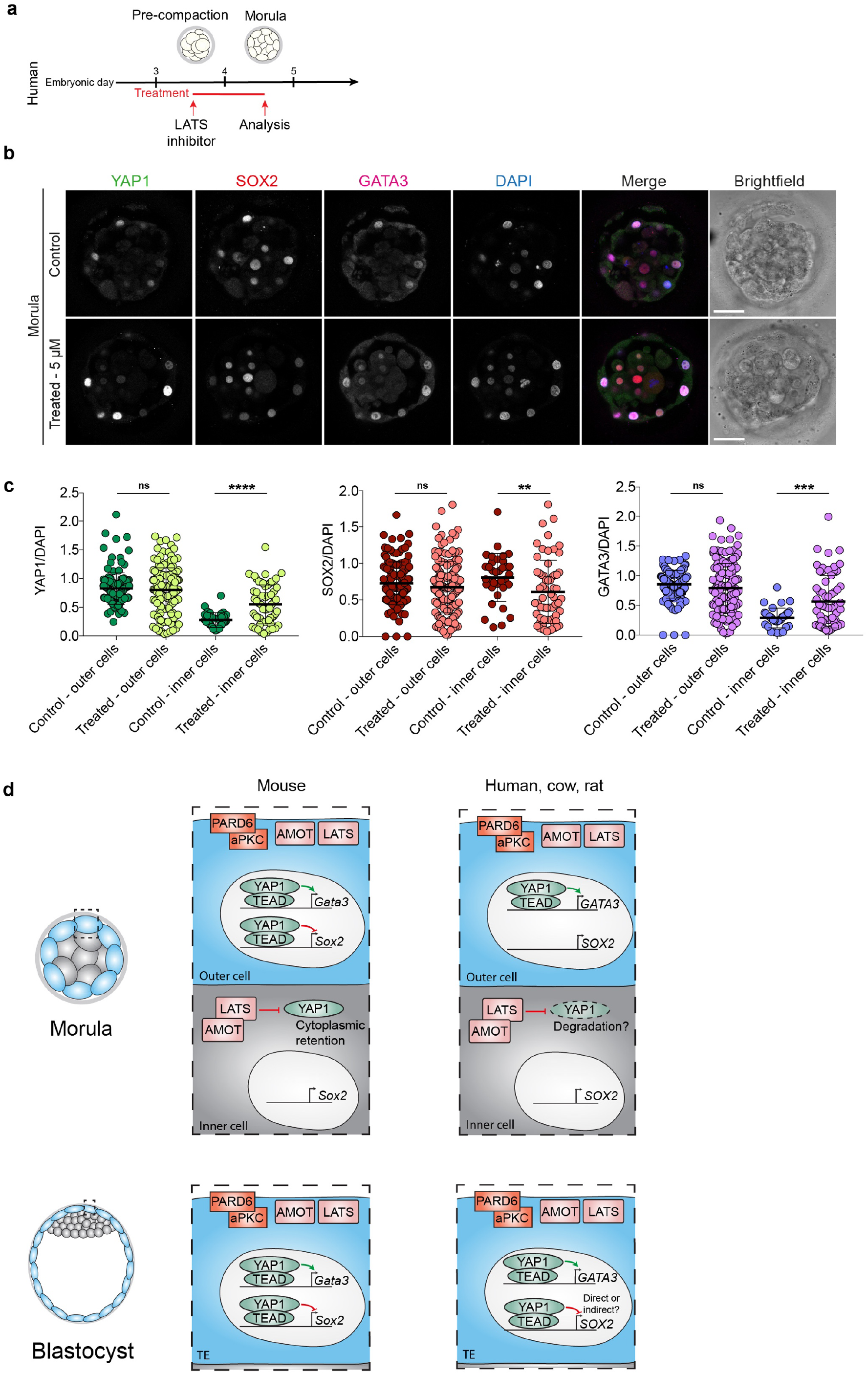
LATS inhibitor treatment in human embryos. **a.** Schematic of LATS inhibitor treatment in human embryos. **b.** Immunofluorescence analysis of YAP1, SOX2, GATA3 and DAPI nuclear staining in control and LATS-inhibitor-treated human morula stage embryos. **c.** Quantification of YAP1, SOX2, GATA3 normalized fluorescence intensity in outer and inner cells in control and LATS-inhibitor-treated human morula stage embryos. (*n* = 394 cells from 11 embryos). Data are presented as mean ± s.d. Two-tailed Mann–Whitney *U* test, ns = not significant, \**P* < 0.05, \*\**P* < 0.01, \*\*\*\**P* < 0.0001. Scale bars, 30 um. **d.** Model for similarities and differences in TE initiation and specification across species. TE = trophectoderm.

Our findings suggest that mouse, rat, cow, and human embryos show an evolutionary conservation of the Hippo pathway, through LATS kinases, despite some notable differences in the timing and expression pattern of differentiation factors downstream of the pathway, consistent with our previous publication (Gerri et al., 2020) (**Figure 5d**). Interestingly, the TE lineage-associated factor CDX2 is expressed in the mouse morula, and detectable in human and cow embryos only at later stages of development (Goissis and Cibelli, 2014a, Niakan and Eggan, 2013). This suggests that despite the conserved role we observed for aPKC-mediated cell polarity, Hippo signaling and initiation of a TE program, not all factors that are known to be required for TE initiation or maintenance are conserved in their expression dynamics and this might reflect differences in the timing or mechanism of lineage specification that necessitates further investigation. Consistent with this, blastocyst chimera and re-aggregation experiments suggest that the timing of irreversible commitment of the TE occurs earlier in the mouse compared to human embryos (De Paepe et al., 2013, Posfai et al., 2017). Further comparative analysis of putative developmental regulators across more species and functional studies of upstream and downstream regulators of early lineage specification would be informative.

The more restricted expression of SOX2 in mouse morula stage embryos compared to the broader expression throughout the morula in rat, cow and human embryos warrants further investigation. In the mouse *Sox2*-null mouse embryos die soon after implantation and exhibit abnormal ICM development (Avilion et al., 2003). Meanwhile, SOX2 is dispensable for initiation of OCT4 and NANOG expression in mouse blastocysts (Wicklow et al., 2014). Interestingly, SOX2 is required for OCT4 and NANOG expression in cow embryos (Luo et al., 2021). Moreover, human SOX2 knockdown studies suggest a requirement for embryo genome activation (Gao et al., 2018). The YAP1/TEAD axis has been shown to function directly upstream of SOX2 in the mouse, but it is unclear whether YAP1 and TEAD4 function similarly in the other species (Frum et al., 2018, Frum et al., 2019). The differences we observed in the detection of cytoplasmic YAP1 at the morula stage is consistent with morphokinetics analysis whereby mouse morula cells undergo more rounds of cell division with the ability of inside and outside cells to switch position and fate (Watanabe et al., 2014, Posfai et al., 2017). This is because our studies suggest that the degradation of YAP1 and rapid exit from the initiation to end of compaction may be a more evolutionally conserved mechanism. It will be interesting to determine through lineage tracing and live imaging analysis if this is associated with cells undergoing fewer fate changes at the morula stage once they are positioned as inside or outside cells. This would be consistent with ICM and TE cells being retained in the same proportion as inner and outer cells at the morula stage in human and cow embryos (Gerri et al., 2020). This, together with the comparatively more rapid changes in SOX2 and GATA3 expression following LATS kinase inhibition in the mouse, suggests molecular differences in regulation of these factors and/or temporal differences in protein turnover across mammals, as suggested for other biological processes (Rayon et al., 2020, Matsuda et al., 2020). In the future, it will be interesting to investigate whether the prolonged expression of SOX2 in TE cells in cow and human blastocysts is mechanistically linked to the late expression of CDX2, as these two transcription factors, considered to be master regulators of early lineage specification, might possess species-specific functions in a fundamental developmental process.

Several molecular differences between mouse and other mammalian species have been reported over recent years, yet it remained unclear if these were rodent- or mouse-specific characteristics. Our data show that some mouse-specific phenotypes are indeed not shared with the rat, which appears more similar to cow and human embryos. In conclusion, our results highlight the importance of cross-species comparisons to elucidate conserved and divergent developmental mechanisms and highlight the importance of the mouse in uncovering fundamental principles that can be investigated in other species.

## Acknowledgements

We thank the donors whose contributions have enabled this research; M. Wilding, H. Premannandan and A. Srikantharajah for the coordination and donation of embryos to our research project; N. Tapon Lab for suggesting us the LATS inhibitor used in this work; members of the laboratories of K.K.N., J. M. A. Turner, R. Lovell-Badge, J. Briscoe, as well as M. Marass, N. Tapon, and A.M. Romao for advice and feedback on the manuscript; the Advanced Light Microscopy and Biological Research Facilities (Francis Crick Institute). Work in the laboratory of H.V.d.V. was funded by the Fonds Wetenschappelijk Onderzoek Flanders (FWO G075222) and the Wetenschappelijk Fonds Willy Gepts (WFWG, UZ Brussel, G142). Work in the laboratory of A.A.F.-N. was supported by Comparative Biomedical Sciences Departmental fund from the Royal Veterinary College. Work in the laboratory of K.K.N. was supported by the Wellcome Trust (221856/Z/20/Z). Work in the laboratory of K.K.N. was also supported by the Francis Crick Institute, which receives its core funding from Cancer Research UK (FC001120), the UK Medical Research Council (FC001120) and the Wellcome Trust (FC001120). For the purpose of Open Access, the authors have applied a CC BY public copyright licence to any Author Accepted Manuscript version arising from this submission.

## Author contributions

C.G. and K.K.N. conceived the study; C.G. and K.K.N. designed the experiments; C.G., K.K.N., A.M., G.M.S., S.B., and C.S.S. performed experiments; G.M.S. and M.R. performed experiments on human embryos in Brussels; J.L., C.M., C.H., S.H., K.E., P.S., and L.C. coordinated the donation of embryos to the research project in London; H.V.d.V. supervised experiments on human embryos in Brussels; C.G., K.K.N., G.M.S., S.B., M.R., and H.V.d.V. analyzed data; A.A.F.-N., D.H., and G.M.S. provided cow ovaries; A.A.F.-N. supervised experiments on cow embryos; C.G. wrote the manuscript with help from all of the authors.

## Competing interests

The authors declare no competing interests.

## Figure supplement legends

**Figure supplement 1.**
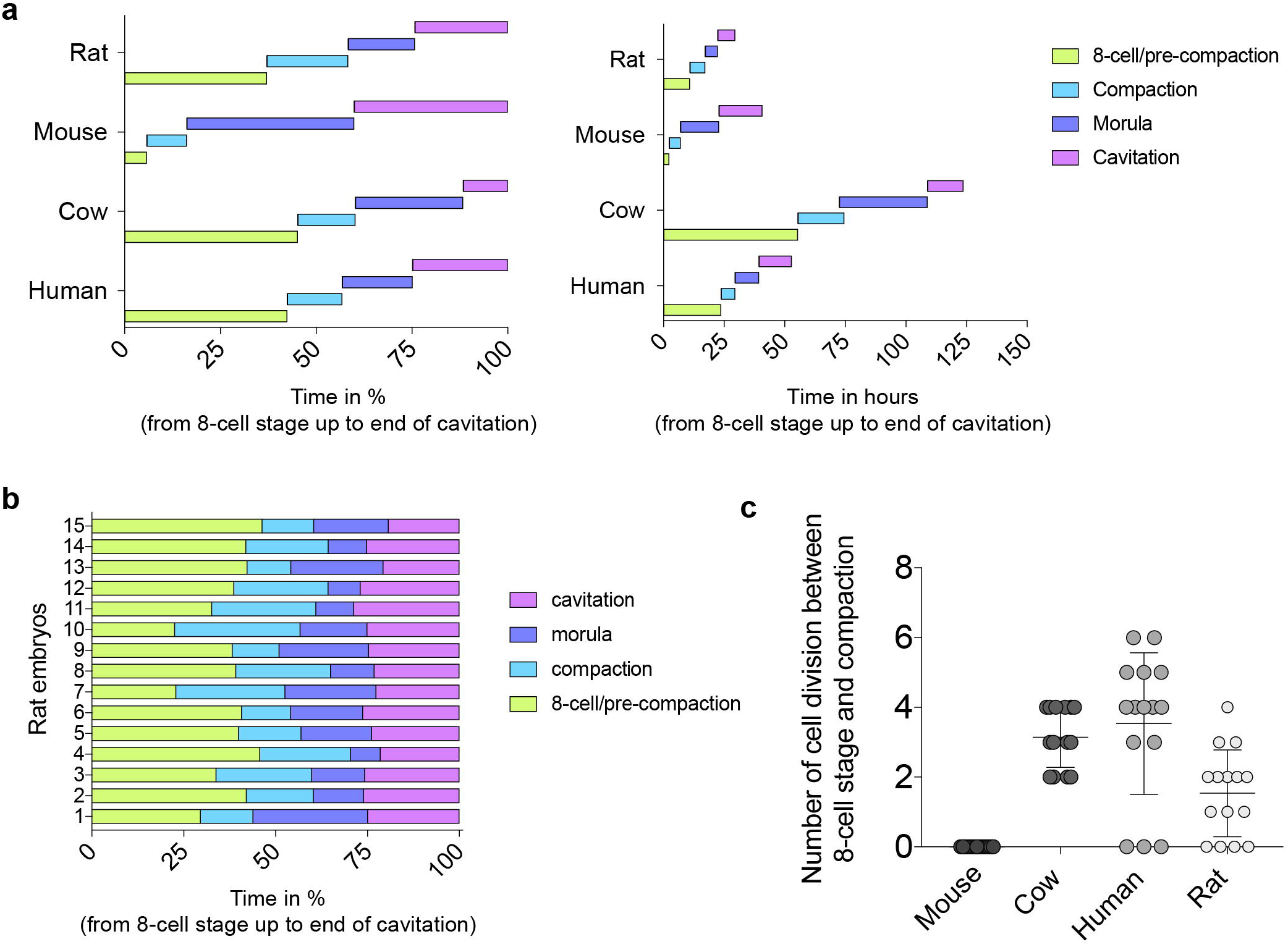
Morphokinetic analysis. **a.** Morphokinetic analysis of rat, mouse, cow and human pre-implantation development, showing relative time in percentage and in hours (from 8-cell stage to end of cavitation). n = 15 for rat, mouse and cow embryos, and n = 16 for human embryos. **b.** Morphokinetic analysis of each of the rat pre-implantation embryos used in the analysis, showing relative time in percentage (from the 8-cell stage up to the end of cavitation). **c.** Quantification of the number of individual cells that divided between 8-cell stage and start of compaction in rat, mouse, cow and human embryos. n =15 embryos for rat, mouse and human, n = 14 embryo for cow. The data shown for mouse, cow and human embryos have already been reported in Gerri et al., Nature 2020. We are reporting these data again to compare them to the newly developed data of rat embryos.

**Figure supplement 2.**
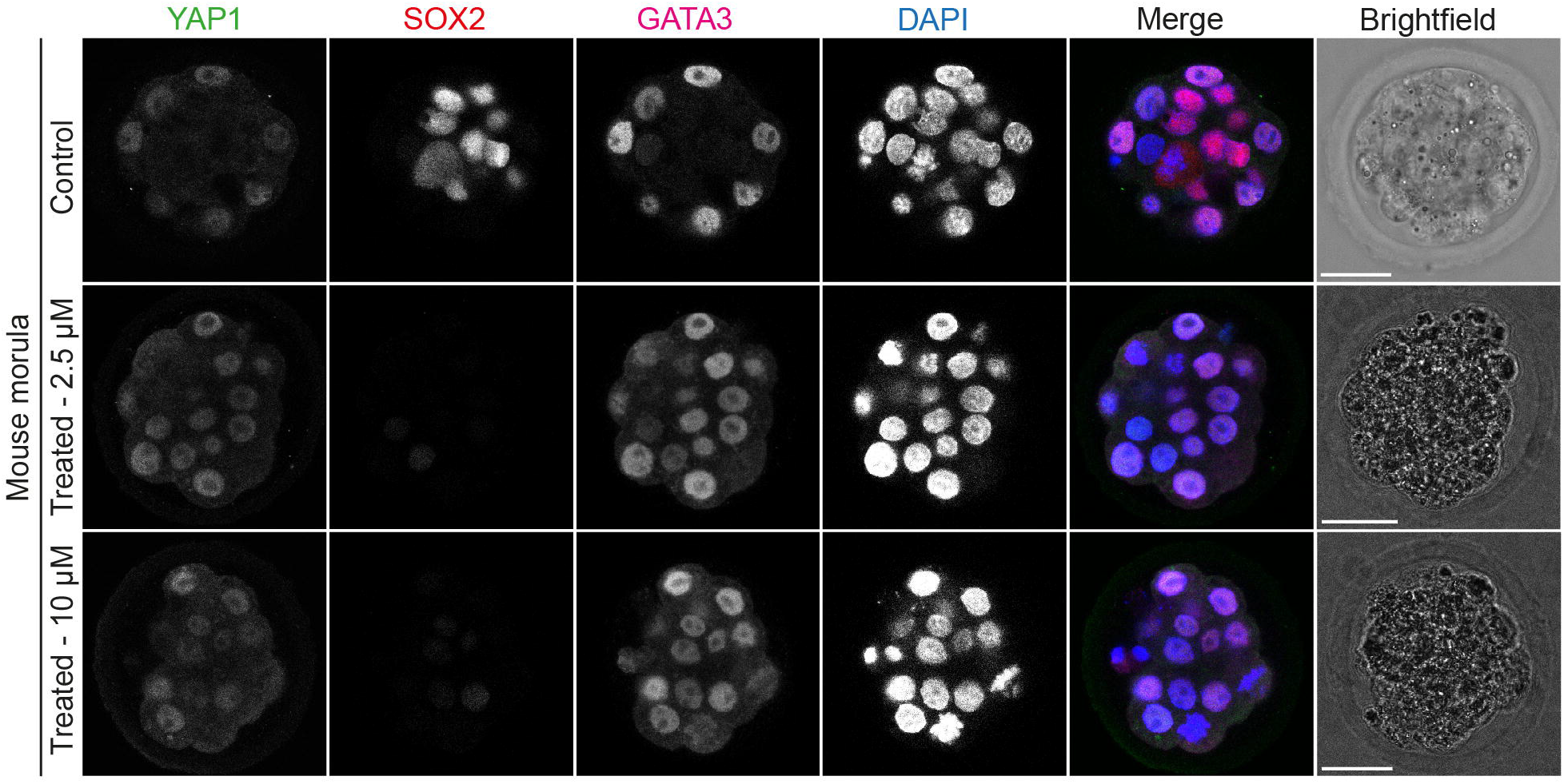
LATS inhibitor dose response experiment in mouse embryos. Immunofluorescence analysis of YAP1, SOX2, GATA3 and DAPI nuclear staining in mouse morula embryos treated with different concentrations of LATS inhibitor (*n* = reported in Table supplement 1). Scale bars, 30 um.

**Figure supplement 3.**
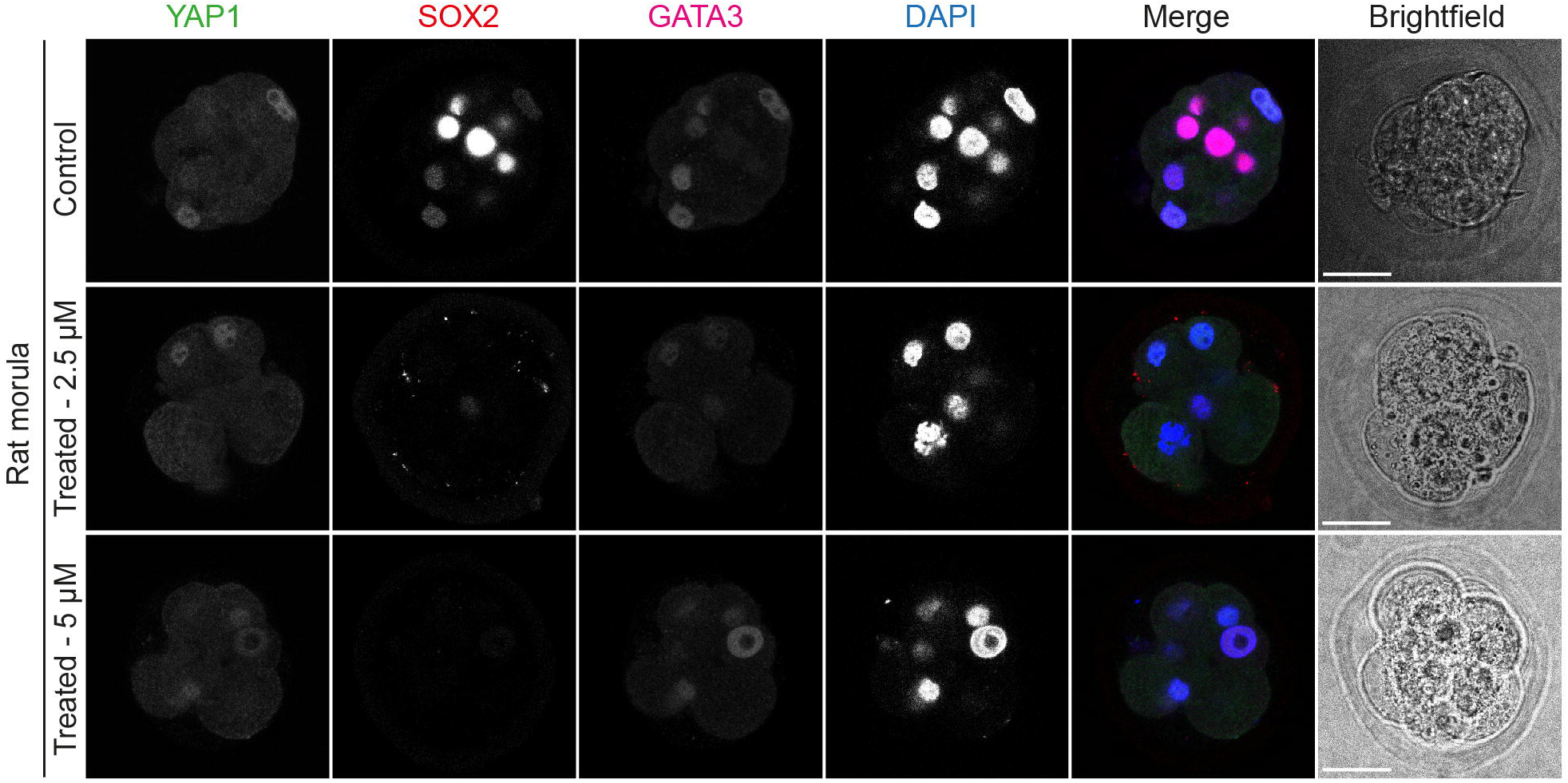
LATS inhibitor dose response experiment in rat embryos. Immunofluorescence analysis of YAP1, SOX2, GATA3 and DAPI nuclear staining in rat morula embryos treated with different concentrations of LATS inhibitor (*n* = reported in Table supplement 2). Scale bars, 30 um.

**Figure supplement 4.**
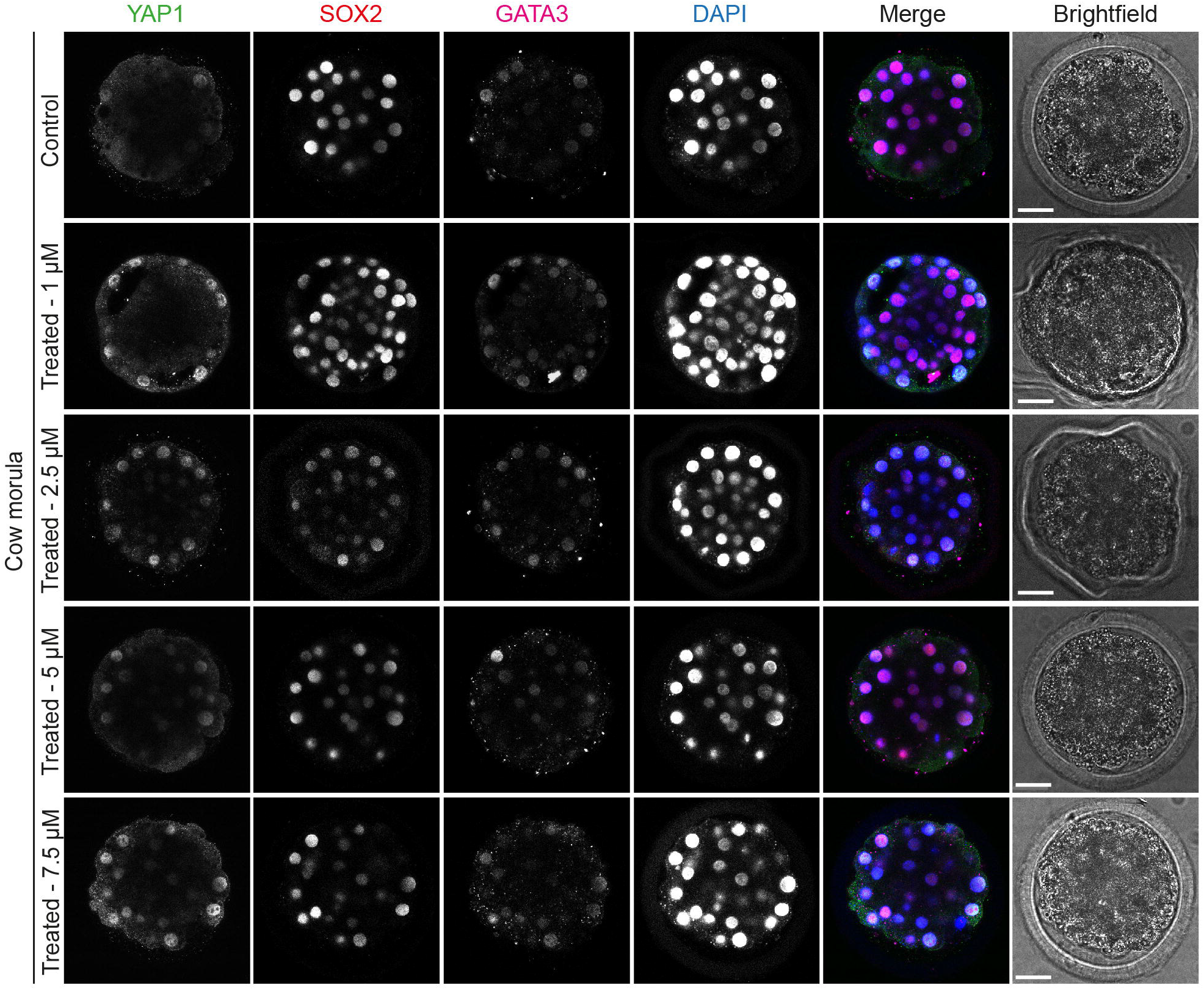
LATS inhibitor dose response experiment in cow embryos. Immunofluorescence analysis of YAP1, SOX2, GATA3 and DAPI nuclear staining in cow morula embryos treated with different concentrations of LATS inhibitor (*n* = reported in Table supplement 3). Scale bars, 30 um.

**Figure supplement 5.**
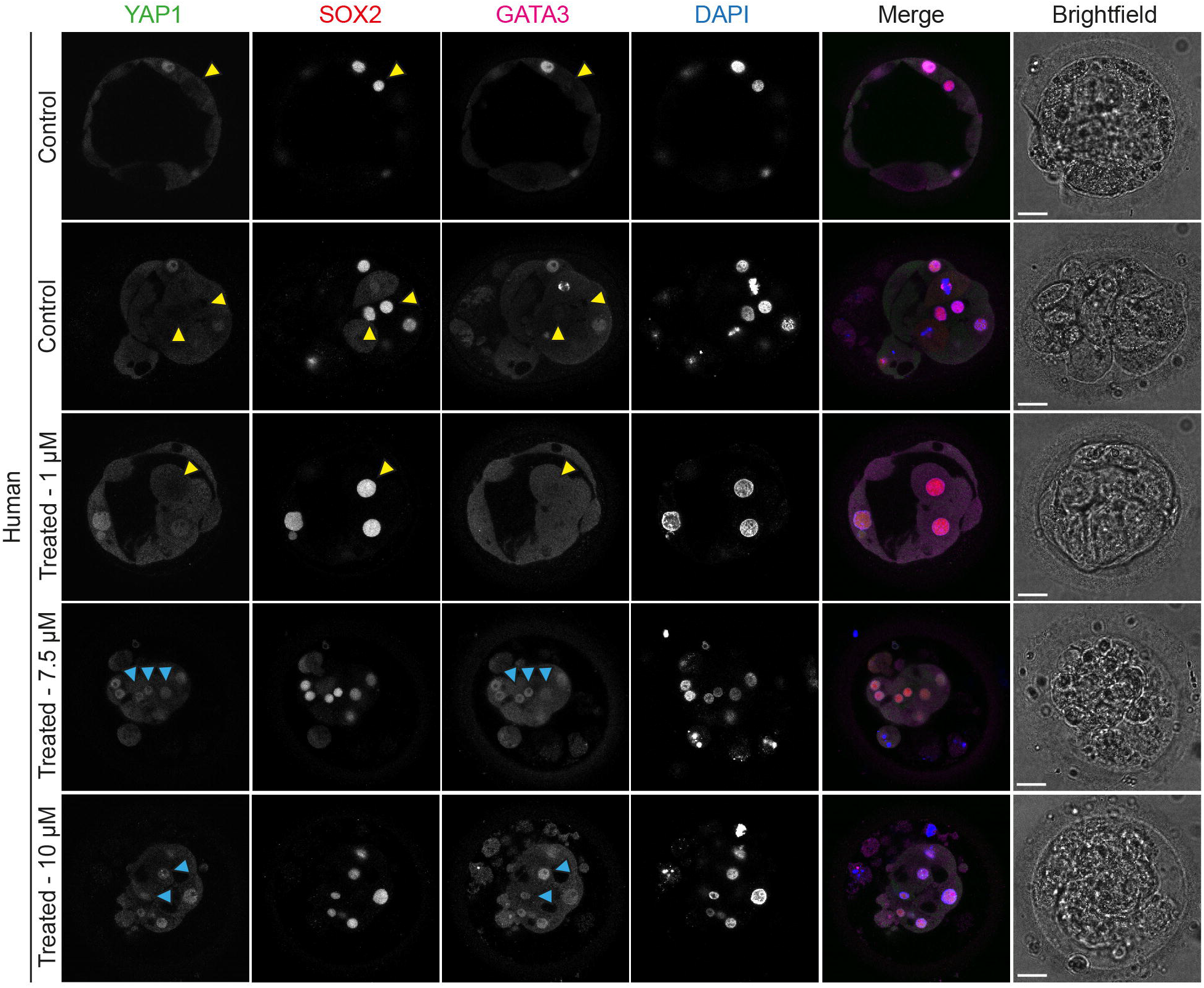
LATS inhibitor dose response experiment in human embryos. Immunofluorescence analysis of YAP1, SOX2, GATA3 and DAPI nuclear staining in human morula embryos treated with different concentrations of LATS inhibitor (*n* = reported in Table supplement 4). Yellow arrowheads point inner/ICM cells expressing only SOX2. Cyan arrowheads point to inner cells expressing SOX2, YAP1 and GATA3. Scale bars, 30 um.

## Table supplement legends

**Table supplement 1. LATS inhibitor concentrations tested in mouse embryos.**

List of concentrations of the LATS inhibitor tested in mouse embryos, treating from the 4-cell to morula stage or to blastocyst stage. We report the total number of embryos treated, the number of embryos arrested during the treatment and number of embryos that reached the analyzed stage.

**Table supplement 2. LATS inhibitor concentrations tested in rat embryos.**

List of concentrations of the LATS inhibitor tested in rat embryos, treating from the 4-cell to morula stage or to blastocyst stage. We report the total number of embryos treated, the number of embryos arrested during the treatment and number of embryos that reached the analyzed stage.

**Table supplement 3. LATS inhibitor concentrations tested in cow embryos.**

List of concentrations of the LATS inhibitor tested in cow embryos, treating from pre-compaction to morula stage or to blastocyst stage. We report the total number of embryos treated, the number of embryos arrested during the treatment and number of embryos that reached the analyzed stage.

**Table supplement 4. LATS inhibitor concentrations tested in human embryos.**

List of concentrations of the LATS inhibitor tested in human embryos, treating from pre-compaction to morula stage. We report the total number of embryos treated, the number of embryos arrested during the treatment and number of embryos that reached the morula stage.

**Table supplement 5. Antibodies used in this study.**

List of antibodies used in this study.

## Video legends

**Video 1.**

Mouse pre-implantation development. Time-lapse Embryoscope+ video of a mouse pronuclear stage zygote developing to the blastocyst stage.

**Video 2.**

Rat pre-implantation development. Time-lapse Embryoscope+ video of a rat 2-cell stage embryo developing to the blastocyst stage.

**Video 3.**

Cow pre-implantation development. Time-lapse Embryoscope video of a cow 2-cell stage embryo developing to the blastocyst stage.

**Video 4.**

Human pre-implantation development. Time-lapse Embryoscope+ video of a human pronuclear stage zygote developing to the blastocyst stage.

## Materials and methods

### Ethics statement

#### Experiments performed in the UK

This study was approved by the UK Human Fertilisation and Embryology Authority (HFEA): research licence number 0162, and the Health Research Authority’s Research Ethics Committee (Cambridge Central reference number 19/EE/0297).

The process of licence approval entailed independent peer review along with consideration by the HFEA Licence and Executive Committees and the Cambridge Central Research Ethics Committee. Our research is compliant with the HFEA Code of Practice and has undergone inspections by the HFEA since the licence was granted. Research donors were recruited from patients at Bourn Hall clinic, Create Fertility, Hewitt Fertility Centre and Homerton University Hospital.

Informed consent was obtained from all couples that donated spare embryos following IVF treatment. Before giving consent, people donating embryos were provided with all of the necessary information about the research project, an opportunity to receive counselling and the conditions that apply within the licence and the HFEA Code of Practice. Donors were informed that embryos used in the experiments would be stopped before 14 days post-fertilization and that subsequent biochemical and genetic studies would be performed. Informed consent was also obtained from donors for all the results of these studies to be published in scientific journals. No financial inducements were offered for donation. Consent was not obtained to perform genetic tests on patients and no such tests were performed. The patient information sheets and consent document provided to patients are publicly available (https://www.crick.ac.uk/research/a-z-researchers/researchers-k-o/kathy-niakan/hfea-licence/). Embryos surplus to the patient’s IVF treatment were donated cryopreserved and were transferred to the Francis Crick Institute where they were thawed and used in the research project.

#### Experiments performed in Belgium

The use of human embryos donated to research was allowed by the Local Ethical Committee of UZ Brussel (BUN 1432021000526) and the Belgian Federal Committee for research on human embryos (AdV087). The embryos were surplus after IVF treatment in Brussels IVF at UZ Brussel. The embryos were cryopreserved and donated to research following informed consent and after the legally determined period of five years of cryopreservation.

#### Human embryo thaw

For embryos thawed in the UK, slow frozen human cleavage stage embryos were thawed using Quinn’s Advantage thaw kit (Origio; ART-8016). Briefly, with Quinn’s Advantage thaw kit, after thawing the embryos were transferred to 0.5% sucrose thawing medium and incubated for 5 min at 37 °C, followed by 0.2% sucrose thawing medium for 10 min at 37 °C. The embryos were then washed through seven drops of diluent solution before culture. For vitrified cleavage stage embryos were thawed using vitrification thaw kit (Irvine Scientific; 90137-SO). With vitrification, embryos were thawed in TS thawing solution for 1 minute and transferred to DS thawing solution for 4 min at RT. Then, embryos were washed twice in washing solution for 4 min at RT before culture. For embryos donated in Belgium, the embryos were thawed as previously described (De Paepe et al., 2019).

#### Mouse zygote collection

All animal research was performed in accordance with the UK Home Office regulations under project licence PP8826065, which passed ethical review by the Francis Crick Institute Animal Welfare Review Board in 2019. Within the Biological Research Facility (BRF) animal units of the Francis Crick institute, mice had *ad libitum* access to feed and water; and were housed in individually ventilated cages maintained at 22°C with 60% humidity on a 12 hour light-dark cycles. Three- to -four to eight-week-old (C57BL6 × CBA) F1 female mice were super-ovulated using injection of 5 IU of pregnant mare serum gonadotrophin (PMSG; Sigma-Aldrich). Forty-eight hours after PMSG injection, 5 IU of human chorionic gonadotrophin (HCG; Sigma-Aldrich) was administered. Superovulated females were set up for mating with eight-week-old or older (C57BL6 × CBA) F1 males. Mouse zygotes were isolated in FHM under mineral oil (Origio; ART-4008-5P) and cumulus cells were removed with hyaluronidase (Sigma-Aldrich; H4272).

#### Human and mouse embryo culture

Mouse embryos or human embryos were cultured in drops of pre-equilibrated Global medium (LifeGlobal; LGGG-20) supplemented with 10% human serum albumim (LifeGlobal; GHSA-125) and overlaid with mineral oil (Origio; ART-4008-5P). Preimplantation embryos were incubated at 37 °C and 5% CO_2_ in an EmbryoScope+ time-lapse incubator (Vitrolife) and cultured up to the day of analysis. For embryos donated in Belgium, the embryos were thawed as previously described (De Paepe et al., 2019).

#### Rat embryo collection and culture

Superovulated and mated 7-9 week old Sprague Dawley females were purchased by Charles River UK Limited. Rat 2-cell embryos were isolated in FHM and then cultured in drops of pre-equilibrated rat embryo culture medium (mR1ECM, Cosmo Bio Co. Ltd., CSR-R-N174) and overlaid with mineral oil (Origio; ART-4008-5P). mbryos were incubated at 37 °C and 5% CO_2_ and cultured up to the day of analysis.

#### Cow embryo generation and culture

Cow ovaries were obtained from the abattoir. Frozen bull sperm was obtained from UK Sire Services. Oocyte isolation, culture, and fertilization were performed using the IVF Bioscience media suite and the manufacturer protocol, with small changes. In brief, cumulus-oocyte complexes (COCs) were aspirated from antral follicles using an 18-gauge needle mounted on a 5 mL disposable syringe. The aspirated follicular fluid was transferred to a 50 mL falcon tube kept at 38.5C. Only fully-grown oocytes with a homogeneous cytoplasm and at least 3-5 complete layers of compact cumulus cells were selected for the experiments. COCs were washed and transferred to pre-warmed and equilibrated BO-IVM media (IVF Biosciences) and incubated at 38.8°C and 5% CO_2_. After 20 hours, frozen semen was thawed and transferred in 4 mL of pre-warmed BO-Semen media (IVF Bioscience) and centrifuged for 5 min at 328*g*. The pellet was resuspended in 4 mL of pre-warmed BO-Semen media and centrifuged for 5 min at 328*g*. The supernatant was discarded and sperm cells were gently resuspended in 1 ml of BO-IVF (IVF Bioscience) and counted using a Bürker chamber. The semen was diluted with BO-IVF (IVF Bioscience) to a concentration of 1×10^6^ spermatozoa and 500 ul wells of diluted semen were prepared. COCs were partially denuded using a P200 and washed in BO-WASH media (IVF Biosciences). COCs were washed into pre-warmed and equilibrated BO-IVF media (IVF Biosciences) and then placed in the wells with the diluted semen. Insemination was performed at 38.5°C, 5% CO_2_ for 20 hours. Zygotes were retrieved after insemination and gently denuded of cumulus cells in pre-warmed BO-WASH medium (IVF bioscience). Zygotes were washed into pre-warmed and equilibrated BO-IVC media (IVF Biosciences) and incubated at 38.5°C, 5% CO_2_ and 5% O_2_.

#### Morphokinetic analysis and embryo staging

Time-lapse imaging was performed using an EmbryoScope+. 8-cell stage was considered to begin when the embryos display 8 obvious blastomeres. The beginning of compaction was defined when blastomeres start to flatten and adhere to each other. The morula stage starts when the embryos appear as a compacted group of cells until the formation of small microlumens. We considered the blastocyst stage to start when embryos show a single dominant blastocoel cavity. In our analysis, cavitation was considered from the end of the morula stage until the formation of an expanded blastocyst, when TE cells touch and start to stretch the zona pellucida, causing zona pellucida thinning. Hatched blastocyst was defined as when part of the blastocyst was hatched outside of the zona pellucida. In addition, from time-lapse imaging we manually counted the number of cell divisions occurring between the 8-cell stage and beginning of compaction. Data on mouse, cow and human embryos are already reported in our previous publication (Gerri et al Nature 2020). We used these data to compared them to our new analysis on rat embryos.

#### Immunofluorescence

Due to high background signal in the zona pellucida of cow embryos, prior to fixation the zona pellucida was removed with acidic Tyrode (Sigma) with 1 mg/ml polyvinyl alcohol (Sigma). Embryos were fixed with freshly prepared 4% paraformaldehyde in PBS that was pre-chilled at 4°C. Embryo fixation was performed for 20 min at RT and then the embryos were transferred through 3 washes of 1X PBS with 0.1% Tween-20 to remove residual paraformaldehyde. Embryos were permeabilized with 1X PBS with 0.5% Triton X-100 and then blocked in blocking solution (3% BSA in 1X PBS with 0.2% Triton X-100) for 2 h at RT on a rotating shaker. Then, embryos were incubated with primary antibodies diluted in blocking solution overnight at 4°C on rotating shaker. The antibodies and the concentrations used are reported in Supplementary Table 11. The following day, embryos were washed in 1X PBS with 0.2% Triton X-100 for 20 min at RT on a rotating shaker and then incubated with secondary antibodies diluted in blocking solution for 1 h at RT on a rotating shaker in the dark. Next, embryos were washed in 1X PBS with 0.2% Triton X-100 for 20 min at RT on rotating shaker. Finally, embryos were placed in 1X PBS with 0.1% Tween-20 with Vectashield and DAPI mounting medium (Vector Lab; H-1200) (1:30 dilution). Phalloidin staining was performed after secondary antibody incubation diluted in blocking solution for 1 h at RT on a rotating shaker in the dark. Embryos were placed on μ-Slide 8 well dishes (Ibidi; 80826) for confocal imaging.

#### Quantification of YAP1, GATA3, SOX2 expression

For morula stage embryos, the measurement of YAP1, SOX2, and GATA3 expression was performed using the 3D Fiji/ImageJ Suite. Briefly, the nuclei of immunofluorescently stained embryos were segmented automatically based on DAPI or HOECHST signal using Fiji, and YAP1, GATA3, SOX2 and TEAD4 fluorescence intensities were measured within the nuclei. We manually verified that the 3D segmentation did not count the same nucleus twice or fused together two adjacent nuclei, and in that case the measurement was excluded from the quantification.

For blastocyst stage embryos, the quantification required a more sophisticated quantification method as for the high number of small cells in blastocyst, which are very closer to each other. Briefly, first StarDist Fiji plugin was used to segments each cell and create a mask for each image. Then, CellProfiler used the mask files from StarDist and measure intensity of each cell, in each channel and in respect to the DAPI channel. These objects where then filtered to remove any objects that are touching the edge of the image. Objects’ size and shape are then measured. After which, objects are then filtered again; this time to exclude any objects that are less than 300px^2, or over 6000px^2 (as pre-determined parameters after some trial and error). This filtering therefore excludes any potential errors in the segmentation by StarDist; such as any cells that had fused together, dividing cells (nuclear DNA fragments), or other artefacts.

Individual objects are then tracked through the z stack, so all z stack images can correspond with an individual blastomere. The final object intensity for each channel is then measured. In addition, we manually verified that the 3D segmentation did not count the same nucleus twice or fused together two adjacent nuclei, and in that case the measurement was excluded from the quantification.

#### Quantification of aPKC expression at the apical membrane

The fluorescence intensity profile of aPKC and AMOT at the apical domain of outer blastomeres were determined using the plot profile tool in Fiji. We draw a ~1-μm-thick line on the apical domain and measure the mean fluorescence intensity of aPKC. We pick confocal slices cutting the cell in the middle, in order to cover the whole apical domain, as previously performed.

#### Confocal Imaging

Confocal immunofluorescence images were taken with a SP8 confocal microscopes and 63x glycerol objective and 1-2-μm-thick optical sections were collected. For embryos imaged in Belgium, images were taken with a LSM800 (Zeiss) confocal microscope.

#### Inhibitor treatment

The LATS inhibitor (Z730688380, Enamine) was dissolved in DMSO to 100 mM stock concentration and diluted at the required concentrations in pre-equilibrated embryo culture media. A dose-response experiment with analysis at the morula stage was performed in all the species. The following concentrations were used for the characterization experiments: 5 μM for mouse embryos, 1 μM for rat embryos, 10 μM for cow embryos and 5 μM for human embryos. The optimal concentration was selected with the dose-response experiment by assessing the effects of the inhibitor on embryo viability (See Tables S1-S4) and phenotypic results based on YAP1, GATA3 and SOX2 expression at the morula stage shown in Figures S2-S5. The optimal concentration was then used to perform the experiments shown in the main Figures 2-5. Control embryos were developed in pre-equilibrated media where the same volume of DMSO was added. It is important to note that for the treatment dishes, we diluted our inhibitor stock in order to add a maximum of 1 μl of inhibitor, and consequently 1 μl of DMSO for the control dishes, diluted in 500 μl of culture medium each.

#### Statistical analysis

All statistical analyses in this study were performed using GraphPad Prism 6.0 or using the R environment for statistical computing. The number of cells or embryos analyzed (*n*), statistical tests, and *p* values are all stated in each figure or figure legend. The *t*-test is an unpaired two-tailed *t*-test and two-tailed Mann Whitney U test. Data are represented as mean ± s.d. Unless otherwise noted, each experiment was performed at least three times.

## Notes

### Competing Interest Statement

The authors have declared no competing interest.

## References

Alarcon, V. B. 2010. Cell polarity regulator PARD6B is essential for trophectoderm formation in the preimplantation mouse embryo. Biol Reprod, 83, 347–58.

Avilion, A. A., Nicolis, S. K., Pevny, L. H., Perez, L., Vivian, N. & Lovell-Badge, R. 2003. Multipotent cell lineages in early mouse development depend on SOX2 function. Genes Dev, 17, 126–40.

Berg, D. K., Smith, C. S., Pearton, D. J., Wells, D. N., Broadhurst, R., Donnison, M. & Pfeffer, P. L. 2011. Trophectoderm lineage determination in cattle. Dev Cell, 20, 244–55.

Cauffman, G., De Rycke, M., Sermon, K., Liebaers, I. & Van de Velde, H. 2009. Markers that define stemness in ESC are unable to identify the totipotent cells in human preimplantation embryos. Hum Reprod, 24, 63–70.

Cockburn, K., Biechele, S., Garner, J. & Rossant, J. 2013. The Hippo pathway member Nf2 is required for inner cell mass specification. Curr Biol, 23, 1195–201.

Cockburn, K. & Rossant, J. 2010. Making the blastocyst: lessons from the mouse. The Journal of Clinical Investigation, 120, 995–1003.

De Paepe, C., Aberkane, A., Dewandre, D., Essahib, W., Sermon, K., Geens, M., Verheyen, G., Tournaye, H. & Van de Velde, H. 2019. BMP4 plays a role in apoptosis during human preimplantation development. Mol Reprod Dev, 86, 53–62.

De Paepe, C., Cauffman, G., Verloes, A., Sterckx, J., Devroey, P., Tournaye, H., Liebaers, I. & Van de Velde, H. 2013. Human trophectoderm cells are not yet committed. Hum Reprod, 28, 740–9.

Dietrich, J. E. & Hiiragi, T. 2007. Stochastic patterning in the mouse pre-implantation embryo. Development, 134, 4219–31.

Ducibella, T. & Anderson, E. 1975. Cell shape and membrane changes in the eight-cell mouse embryo: Prerequisites for morphogenesis of the blastocyst. Developmental Biology, 47, 45–58.

Frum, T., Murphy, T. M. & Ralston, A. 2018. HIPPO signaling resolves embryonic cell fate conflicts during establishment of pluripotency in vivo. Elife, 7.

Frum, T., Watts, J. L. & Ralston, A. 2019. TEAD4, YAP1 and WWTR1 prevent the premature onset of pluripotency prior to the 16-cell stage. Development, 146.

Gao, L., Wu, K., Liu, Z., Yao, X., Yuan, S., Tao, W., Yi, L., Yu, G., Hou, Z., Fan, D., Tian, Y., Liu, J., Chen, Z. J. & Liu, J. 2018. Chromatin Accessibility Landscape in Human Early Embryos and Its Association with Evolution. Cell, 173, 248–259.e15.

Gerri, C., Mccarthy, A., Alanis-Lobato, G., Demtschenko, A., Bruneau, A., Loubersac, S., Fogarty, N. M. E., Hampshire, D., Elder, K., Snell, P., Christie, L., David, L., Van de Velde, H., Fouladi-Nashta, A. A. & Niakan, K. K. 2020. Initiation of a conserved trophectoderm program in human, cow and mouse embryos. Nature, 587, 443–447.

Goissis, M. D. & Cibelli, J. B. 2014a. Functional characterization of CDX2 during bovine preimplantation development in vitro. Mol Reprod Dev, 81, 962–70.

Goissis, M. D. & Cibelli, J. B. 2014b. Functional characterization of SOX2 in bovine preimplantation embryos. Biol Reprod, 90, 30.

Hirate, Y., Hirahara, S., Inoue, K., Suzuki, A., Alarcon, V. B., Akimoto, K., Hirai, T., Hara, T., Adachi, M., Chida, K., Ohno, S., Marikawa, Y., Nakao, K., Shimono, A. & Sasaki, H. 2013. Polarity-dependent distribution of angiomotin localizes Hippo signaling in preimplantation embryos. Curr Biol, 23, 1181–94.

Kastan, N., Gnedeva, K., Alisch, T., Petelski, A. A., Huggins, D. J., Chiaravalli, J., Aharanov, A., Shakked, A., Tzahor, E., Nagiel, A., Segil, N. & Hudspeth, A. J. 2021. Small-molecule inhibition of Lats kinases may promote Yap-dependent proliferation in postmitotic mammalian tissues. Nature Communications, 12, 3100.

Leung, C. Y. & Zernicka-Goetz, M. 2013. Angiomotin prevents pluripotent lineage differentiation in mouse embryos via Hippo pathway-dependent and-independent mechanisms. Nat Commun, 4, 2251.

Luo, L., Shi, Y., Wang, H., Wang, Z., Dang, Y., Li, S., Wang, S. & Zhang, K. 2021. Base editing in bovine embryos reveals a species-specific role of SOX2 in regulation of pluripotency. bioRxiv, 2021.11.10.468023.

Matsuda, M., Hayashi, H., Garcia-Ojalvo, J., Yoshioka-Kobayashi, K., Kageyama, R., Yamanaka, Y., Ikeya, M., Toguchida, J., Alev, C. & Ebisuya, M. 2020. Species-specific segmentation clock periods are due to differential biochemical reaction speeds. Science, 369, 1450–1455.

Niakan, K. K. & Eggan, K. 2013. Analysis of human embryos from zygote to blastocyst reveals distinct gene expression patterns relative to the mouse. Dev Biol, 375, 54–64.

Nishioka, N., Inoue, K., Adachi, K., Kiyonari, H., Ota, M., Ralston, A., Yabuta, N., Hirahara, S., Stephenson, R. O., Ogonuki, N., Makita, R., Kurihara, H., Morin-Kensicki, E. M., Nojima, H., Rossant, J., Nakao, K., Niwa, H. & Sasaki, H. 2009. The Hippo signaling pathway components Lats and Yap pattern Tead4 activity to distinguish mouse trophectoderm from inner cell mass. Dev Cell, 16, 398–410.

Niwa, H., Toyooka, Y., Shimosato, D., Strumpf, D., Takahashi, K., Yagi, R. & Rossant, J. 2005. Interaction between Oct3/4 and Cdx2 determines trophectoderm differentiation. Cell, 123, 917–29.

Ohsugi, M., Hwang, S. Y., Butz, S., Knowles, B. B., Solter, D. & Kemler, R. 1996. Expression and cell membrane localization of catenins during mouse preimplantation development. Dev Dyn, 206, 391–402.

Plusa, B., Frankenberg, S., Chalmers, A., Hadjantonakis, A.-K., Moore, C. A., Papalopulu, N., Papaioannou, V. E., Glover, D. M. & Zernicka-Goetz, M. 2005. Downregulation of Par3 and aPKC function directs cells towards the ICM in the preimplantation mouse embryo. Journal of Cell Science, 118, 505–515.

Posfai, E., Petropoulos, S., De Barros, F. R. O., Schell, J. P., Jurisica, I., Sandberg, R., Lanner, F. & Rossant, J. 2017. Position- and Hippo signaling-dependent plasticity during lineage segregation in the early mouse embryo. Elife, 6.

Ralston, A., Cox, B. J., Nishioka, N., Sasaki, H., Chea, E., Rugg-Gunn, P., Guo, G., Robson, P., Draper, J. S. & Rossant, J. 2010. Gata3 regulates trophoblast development downstream of Tead4 and in parallel to Cdx2. Development, 137, 395–403.

Rayon, T., Stamataki, D., Perez-Carrasco, R., Garcia-Perez, L., Barrington, C., Melchionda, M., Exelby, K., Lazaro, J., Tybulewicz, V. L. J., Fisher, E. M. C. & Briscoe, J. 2020. Speciesspecific pace of development is associated with differences in protein stability. Science, 369.

Strumpf, D., Mao, C. A., Yamanaka, Y., Ralston, A., Chawengsaksophak, K., Beck, F. & Rossant, J. 2005. Cdx2 is required for correct cell fate specification and differentiation of trophectoderm in the mouse blastocyst. Development, 132, 2093–102.

Vinot, S., Le, T., Ohno, S., Pawson, T., Maro, B. & Louvet-Vallée, S. 2005. Asymmetric distribution of PAR proteins in the mouse embryo begins at the 8-cell stage during compaction. Developmental Biology, 282, 307–319.

Watanabe, T., Biggins, J. S., Tannan, N. B. & Srinivas, S. 2014. Limited predictive value of blastomere angle of division in trophectoderm and inner cell mass specification. Development, 141, 2279–88.

Wicklow, E., Blij, S., Frum, T., Hirate, Y., Lang, R. A., Sasaki, H. & Ralston, A. 2014. HIPPO pathway members restrict SOX2 to the inner cell mass where it promotes ICM fates in the mouse blastocyst. PLoS Genet, 10, e1004618.

Zhu, M., Shahbazi, M., Martin, A., Zhang, C., Sozen, B., Borsos, M., Mandelbaum, R. S., Paulson, R. J., Mole, M. A., Esbert, M., Titus, S., Scott, R. T., Campbell, A., Fishel, S., Gradinaru, V., Zhao, H., Wu, K., Chen, Z.-J., Seli, E., De Los Santos, M. J. & Zernicka Goetz, M. 2021. Human embryo polarization requires PLC signaling to mediate trophectoderm specification. eLife, 10, e65068.

