## Supplementary material for "A conserved role of Hippo signaling in initiation of the first lineage specification event across mammals": Table supplement 1

| **LATS inhibitor treatment from 4-cell to morula stage in mouse embryos** | | | |
|---|---|---|---|
| **LATS inhibitor concentration** | **Number of embryos** | **Number (percentage) of arrested embryos** | **Number (percentage) of morula embryos** |
| DMSO control (volume matched) | 50 | 3 (6%) | 47 (94%) |
| 2.5 µM | 25 | 1 (4%) | 24 (96%) |
| 5 µM | 50 | 4 (8%) | 46 (92%) |
| 10 µM | 25 | 10 (40%) | 15 (60%) |

| **LATS inhibitor treatment from 4-cell to blastocyst stage in mouse embryos** | | | |
|---|---|---|---|
| **LATS inhibitor concentration** | **Number of embryos** | **Number (percentage) of arrested embryos** | **Number (percentage) of blastocyst embryos** |
| DMSO control (volume matched) | 50 | 3 (6%) | 47 (94%) |
| 5 µM | 50 | 2 (4%) | 48 (96%) |
