## Supplementary material for "A conserved role of Hippo signaling in initiation of the first lineage specification event across mammals": Table supplement 2

| LATS inhibitor treatment from 4-cell to morula stage in rat embryos | | | |
|---|---|---|---|
| **LATS inhibitor concentration** | **Number of embryos** | **Number (percentage) of arrested embryos** | **Number (percentage) of morula embryos** |
| DMSO (volume matched) | 50 | 5 (10%) | 45 (90%) |
| 1 µM | 50 | 6 (12%) | 44 (88%) |
| 2.5 µM | 25 | 11 (44%) | 14 (56%) |
| 5 µM | 25 | 16 (64%) | 9 (36%) |

| LATS inhibitor treatment from 4-cell to blastocyst stage in rat embryos | | | |
|---|---|---|---|
| **LATS inhibitor concentration** | **Number of embryos** | **Number (percentage) of arrested embryos** | **Number (percentage) of blastocyst embryos** |
| DMSO (volume matched) | 45 | 5 (11%) | 40 (89%) |
| 1 µM | 45 | 3 (7%) | 42 (93%) |
