## Supplementary material for "A conserved role of Hippo signaling in initiation of the first lineage specification event across mammals": Table supplement 3

| LATS inhibitor treatment from pre-compaction to morula stage in cow embryos | | | |
|---|---|---|---|
| **LATS inhibitor concentration** | **Number of embryos** | **Number (percentage) of arrested embryos** | **Number (percentage) of morula embryos** |
| DMSO (volume matched) | 50 | 3 (6%) | 47 (94%) |
| 1 μM | 22 | 2 (9%) | 20 (91%) |
| 2.5 μM | 18 | 1 (6%) | 17 (94%) |
| 5 μM | 45 | 3 (7%) | 42 (93%) |
| 7.5 μM | 23 | 1 (4%) | 22 (96%) |
| 10 μM | 50 | 5 (10%) | 45 (90%) |

| LATS inhibitor treatment from pre-compaction to blastocyst stage in cow embryos | | | |
|---|---|---|---|
| **LATS inhibitor concentration** | **Number of embryos** | **Number (percentage) of arrested embryos** | **Number (percentage) of blastocyst embryos** |
| DMSO (volume matched) | 40 | 2 (5%) | 38 (95%) |
| 10 μM | 40 | 3 (7%) | 37 (93%) |
