## Supplementary material for "A conserved role of Hippo signaling in initiation of the first lineage specification event across mammals": Table supplement 4

| LATS inhibitor treatment from pre-compaction to morula stage in human embryos | | | |
|---|---|---|---|
| **LATS inhibitor concentration** | **Number of embryos** | **Number (percentage) of arrested embryos** | **Number (percentage) of morula embryos** |
| DMSO (volume matched) | 10 | 0 (0%) | 10 (100%) |
| 1 µM | 2 | 0 (0%) | 2 (100%) |
| 5 µM | 8 | 0 (0%) | 8 (100%) |
| 7.5 µM | 4 | 2 (50%) | 2 (50%) |
| 10 µM | 1 | 0 (0%) | 1 (100%) |
