## Supplementary material for "A conserved role of Hippo signaling in initiation of the first lineage specification event across mammals": Table supplement 5

| List of antibodies used in this study | | | |
|---|---|---|---|
| **Antibody** | **Product Number** | **Source** | **Dilution** |
| GATA3 | AF2605 | R&D | 1:100 (for human, mouse and rat) |
| GATA3 | AB199428 | Abcam | 1:100 (for cow) |
| YAP1 | H00010413-M01 | Abnova | 1:50 |
| SOX2 | 14-9811-82 | eBioscience | 1:100 |
